# The evolution of protein domain repertoires: shedding light on the origins of herpesviruses

**DOI:** 10.1101/423525

**Authors:** Anderson F. Brito, John W. Pinney

**Author notes:** **Corresponding author:** John W. Pinney.

## Abstract

Herpesviruses (HVs) have large genomes that can encode thousands of proteins. Apart from amino acid mutations, protein domain acquisitions, duplications and losses are also common modes of evolution. HV domain repertoires differ across species, and only a core set is shared among all viruses, aspect that raises a question: How have HV domain repertoires diverged while keeping some similarities? To answer such question, we used profile HMMs to search for domains in all possible translated ORFs of fully sequenced HV genomes. With at least 274 domains being identified, we built a matrix of domain counts per species, and applied a parsimony method to reconstruct the ancestral states of these domains along the HV phylogeny. It revealed events of domain gain, duplication and loss over more than 400 millions of years, where Alpha-, Beta- and Gammaherpesviruses expanded and condensed their domain repertoires at distinct rates. Most of the acquired domains perform ‘Modulation and Control’, ‘Envelope’ or ‘Auxiliary’ functions, categories that showed high flexibility (number of domains) and redundancy (number of copies). Conversely, few gains and duplications were observed for domains involved in ‘Capsid assembly and structure’, and ‘DNA Replication, recombination and metabolism’. Among the 41 primordial domains encoded by herpesvirus ancestors, 28 are still found in all present-day HVs. Because of their distinct evolutionary strategies, herpesvirus domain repertoires are very specific at the subfamily, genus and species levels. Differences in domain composition may not just explain HV host range and tissue tropism, but also provide hints to the origins of herpesviruses.

---

Viruses are known by their unorthodox modes of evolution and outstanding strategies of immune evasion and cell infection. Their biological origins is a question that has long been debated, with many theories being proposed, with a consensus (Wessner 2010; Nasir and Caetano-Anolles 2015; Harish et al. 2016), probably because there is no single answer for such complex question (Wessner 2010). Talking about large DNA viruses, such as members of the family *Herpesviridae,* answering this question is even harder, specially taking into consideration their genomic complexity.

*Herpesviridae* is the best-characterized family of large double-stranded DNA viruses (Davison 2007; Davison et al. 2009), being divided in three subfamilies: *Alpha-; Beta-,* and; *Gammaherpesvirinae* (Davison et al. 2009). Among the 75 fully sequenced genomes, ranging in size from 109 to 241 Kbp, the total number of ORFs encoded by them varies from 69 to 223 (Brister et al. 2015). A set of at least 41 core genes are shared among all HVs, which play essential roles in viral infection, such as assembly of virion structure, DNA replication, viral entry and egress pathways (Mocarski 2007).

During infections, herpesviruses (HVs) are known to establish latency in host cells, where their genomes are circularized, packed with histones, and copied by the cellular machinery during mitosis (Grinde 2013); and at this stage, genetic elements from hosts can be captured by viruses via horizontal gene transfers (HGTs) (Holzerlandt et al. 2002). Each viral or host protein is composed by one of more domains, functional units that play specific roles and evolve independently from each other (Vogel et al. 2004). The Pfam database (Finn et al.2014), now on its version 31.0, contains information about more than 16700 domain families (Pfam-A), which are defined based on probabilistic models generated from high-quality position-specific amino acid alignments. When implemented, such models can efficiently identify remote homology between sequences belonging to the same domain family. By identifying the domains of all proteins encoded by viral genomes, their species-specific repertoires can be reconstructed, revealing not just their array of encoded domains, but also the set of molecular functions they are capable to perform. Domain repertoires can evolve under three main events: domain gains (Raftery et al. 2000; Holzerlandt et al. 2002), losses (Albalat and Canestro 2016) and duplications (Gao et al. 2017).

In the present study we extracted all possible ORFs from 75 fully sequenced genomes of herpesviruses, and identified at least 274 protein domains encoded by them. By using a time-calibrated phylogenetic tree of viral species, and a matrix of domain counts for each species, a parsimony model was applied to reconstruct the ancestral states (presence/absence) of domains at all internal nodes of the tree, including its root. This analysis allowed us to reconstruct a parsimonious scenario explaining the evolution of HV domain families by events of domain gain, loss and duplication. It revealed that current herpesviral species encode a core set of 28 domains inherited from their ancestors since approximately 400 million years ago. By classifying the HV domains in functional groups, it was possible to determine that domains implied in virus-host interactions were the main elements acquired by herpesviruses, especially in early periods of their evolution. Our results were interpreted in light of the current hypothesis explaining genomic and viral evolution, and shed light on the possible origins of herpesviruses.

## RESULTS

### The current domain repertoire of herpesviruses

To circumvent the limitations of simple sequence comparisons, which fail at identifying homology between distantly related proteins (Mocarski 2007), in this study we applied HMM profiles generated from Pfam alignments to identify domains encoded by fully sequenced herpesviral genomes. To allow a good balance between sensitivity and selectivity, the per-sequence (independent) and per-domain (conditional) Evalues were defined using a conservative cutoff of 0.001, as used in previous studies (Ekman et al. 2007; Moore and Bornberg-Bauer 2012). Such a robust approach allowed us to detect a non-redundant set of at least 274 domains (Supplemental Table S2), which cluster mostly following the phylogeny of HVs (Figure 1). Interestingly, domains from distinct functional groups have shown to be unequally distributed across the HV subfamilies and species (Figure 2).

**Figure 1.**
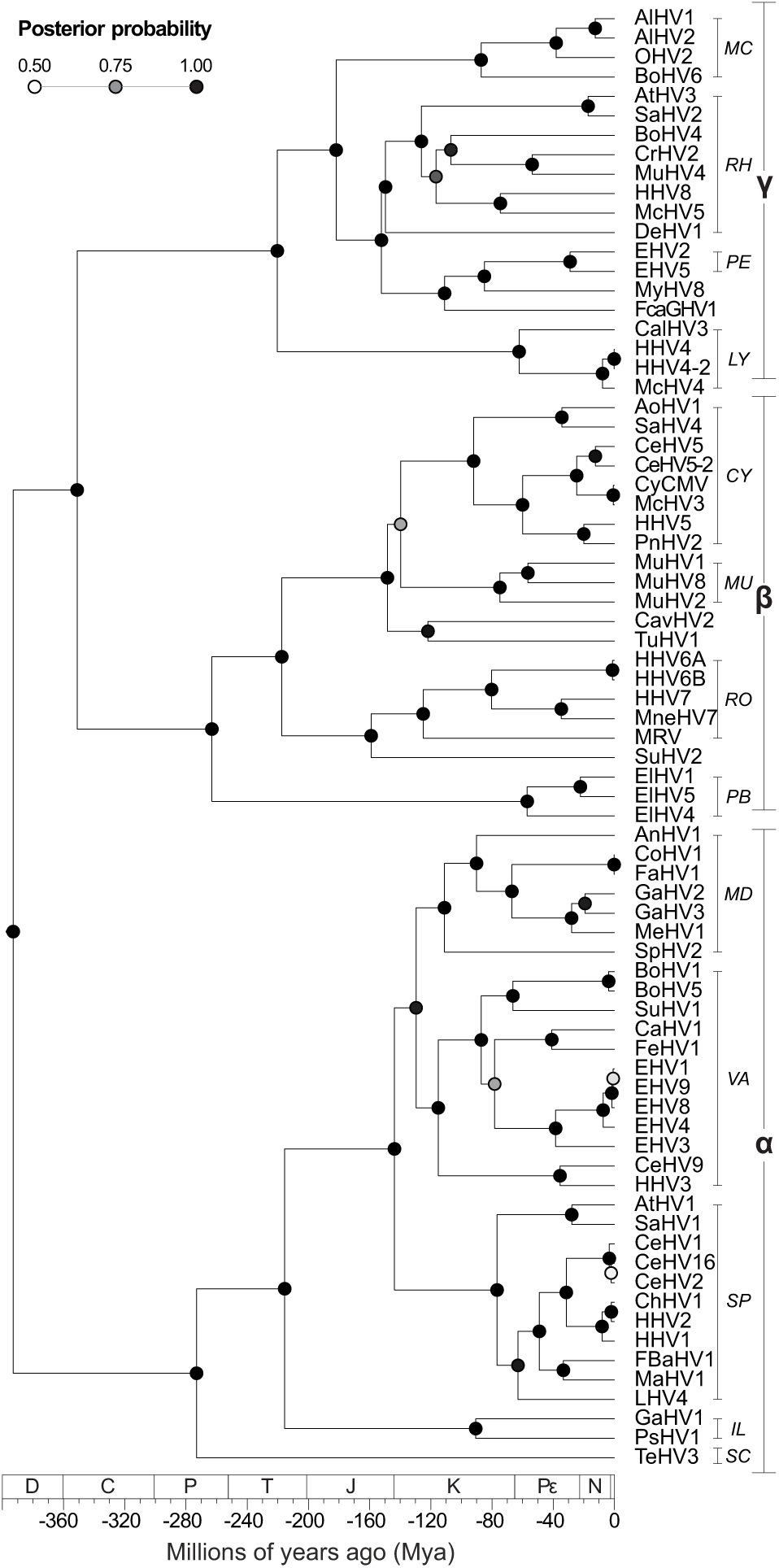
Time-calibrated maximum clade credibility (MCC) tree of fully sequenced herpesviruses. Posterior probabilities are shown at the nodes. Herpesvirus subfamilies are shown as a *(Alphaherpesvirinae);* β *(Betaherpesvirinae)* and y *(Gammaherpesvirinae).* Herpesvirus genera are shown as: CY = *Cytomegalovirus;* IL = *Iltovirus;* LY = *Lymphocryptovirus;* MC = *Macavirus;* MD = *Mardivirus;* MU = *Muromegalovirus;* PB = *Proboscivirus;* PE = *Percavirus;* RH = *Rhadinovirus;* RO = *Roseolovirus;* SC = *Scutavirus;* SP = *Simplexvirus;* and VA = *Varicellovirus.* The geologic time scale is set according to (Gradstein et al. 2012), where D = Devonian period; C = Carboniferous; P = Permian; T = Triassic; J = Jurassic; K = Cretaceous; Pϵ = Paleogene; N = Neogene; and * = Quaternary period (Q).

**Figure 2.**
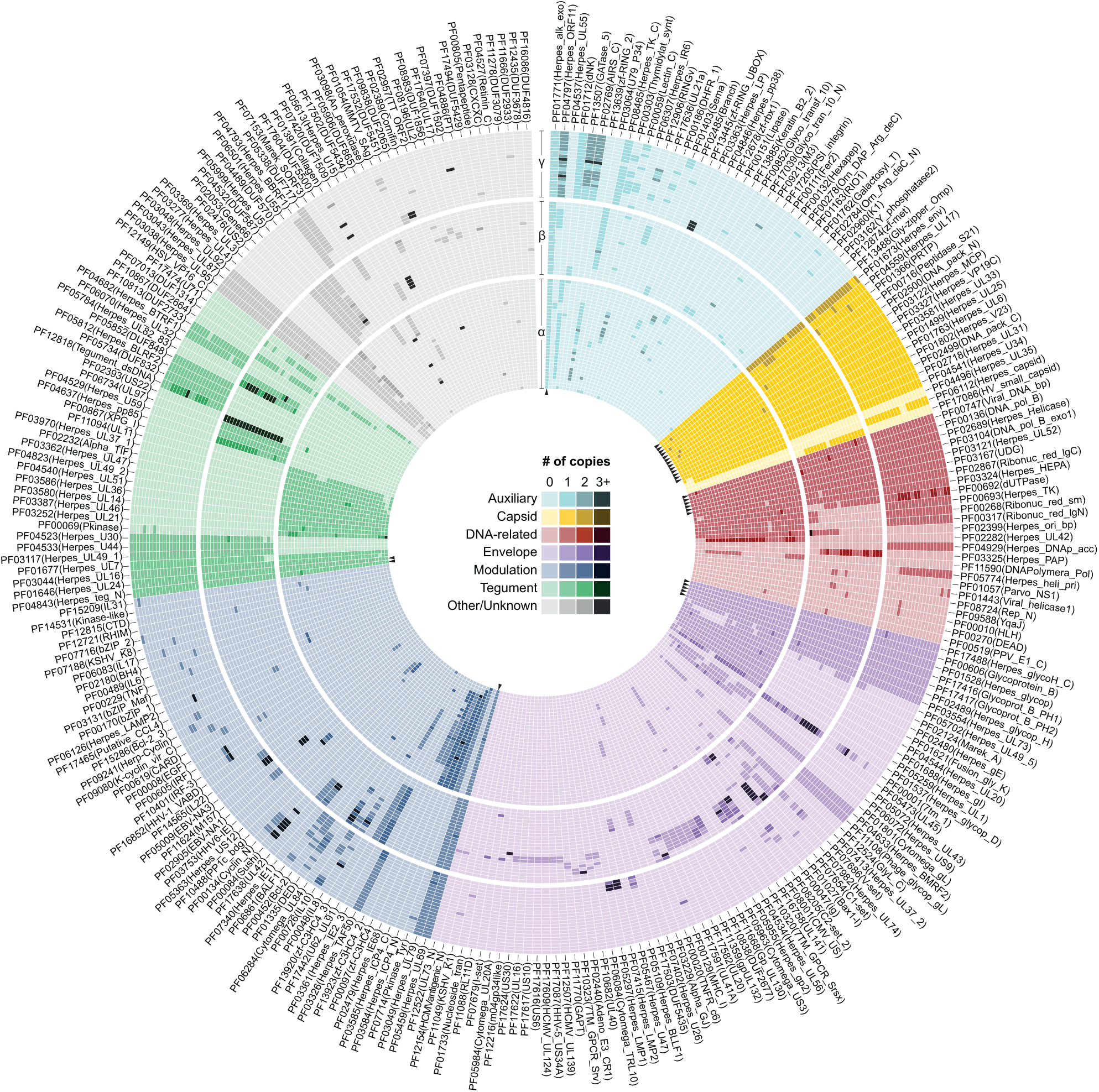
The domain repertoire of herpesviruses. A total of 274 protein domains were identified from ORFs of HV genomes. In this circular map, viral species are positioned following the same vertical order as shown in Figure 1, with heatmaps of distinct subfamilies clearly separated. Domains with similar function were grouped, and the horizontal order of domains (clockwise sense) was defined based on their absolute frequency (i.e. most conserved domains are shown at the first columns of each group). Please observe that the lightest colour shade of each group indicates domains that are absent in some species. As observed, only a small set of domains (n = 28, see arrowheads) was found across all species, and most domains (n = 203) are restricted to a single subfamily. The circular map was designed using Circos (Krzywinski et al. 2009).

A total of 46 domains - most of them involved in ‘Capsid assembly and structure’; ‘Envelope’; and ‘DNA-related processes’ - were found in all subfamilies of herpesviruses (Figure 3). Nevertheless, since some of these domains were lost or have been independently acquired along the evolution, only 28 of them are encoded by all HV species, and as herpesviruses encode on average 75.6 ± 8.4 domains, non-core domains compose nearly two thirds of the HV repertoires.

**Figure 3.**
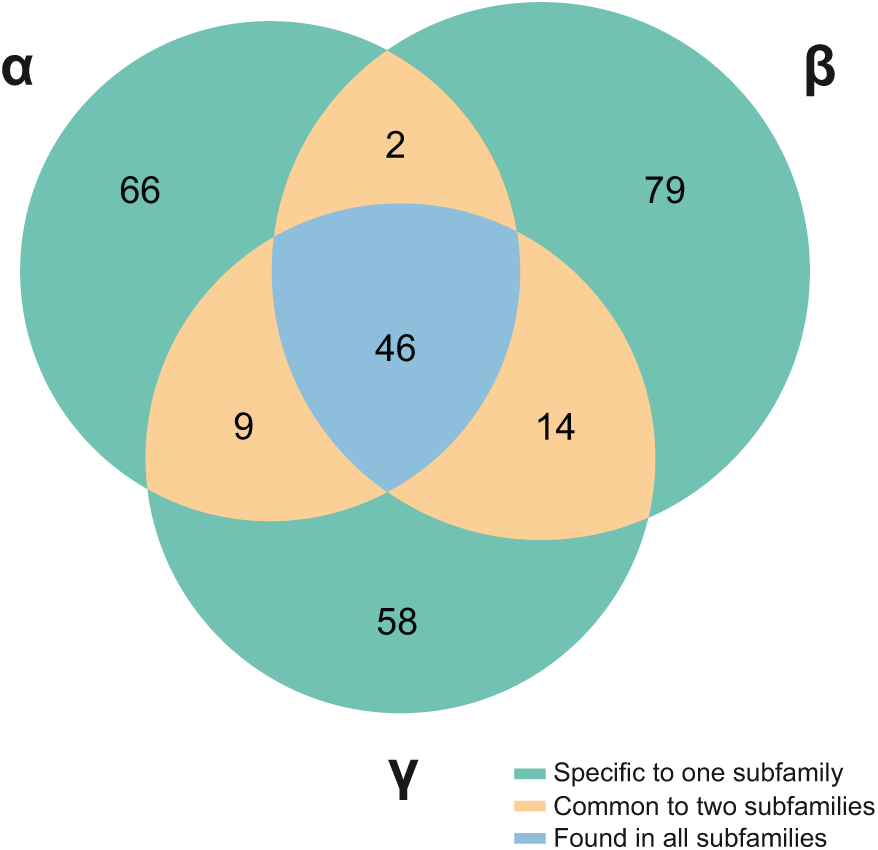
Distribution of protein domains across distinct HV subfamilies (α, β and γ). A set of 46 domains was found in all subfamilies, but not in all species. As highlighted, the large majority of the protein domains of herpesviruses are highly subfamily-specific.

Interestingly, 203 out of 274 domains are specific to a single subfamily, with 73 of them being strict to only one or two HV species, an aspect that highlights a trend for molecular specialization. A few domains are shared between two subfamilies while being absent in a third one, and as expected due to their closer phylogenetic relationship, Beta- and GammaHVs share higher number of domains (n = 14, five of them envelope proteins) (Figure 3). Finally, another group of 46 domains (16.8%) was found to be species-specific, most of them performing auxiliary (enzymatic) functions, or playing a role in the modulation and control of gene expression.

### The interplay among domain gains, losses and duplications

By taking the current set of domains encoded by herpesviral genomes, we reconstructed a possible scenario explaining the origins of herpesvirus domain repertoires. To do that we applied a parsimony reconstruction method that revealed an intense dynamics of domain gains, losses and duplications along more than 400 millions of years of herpesviral evolution (Figure 4). In this study, an event was classified as a ‘gain’ when a new domain, not yet present in the parent of a certain node, was incorporated in the repertoire of a HV lineage. When an existing domain decreased its overall copy number at a child node, a ‘loss’ was recorded, and accordingly, when its frequency increased, it was classified as a ‘duplication’.

**Figure 4.**
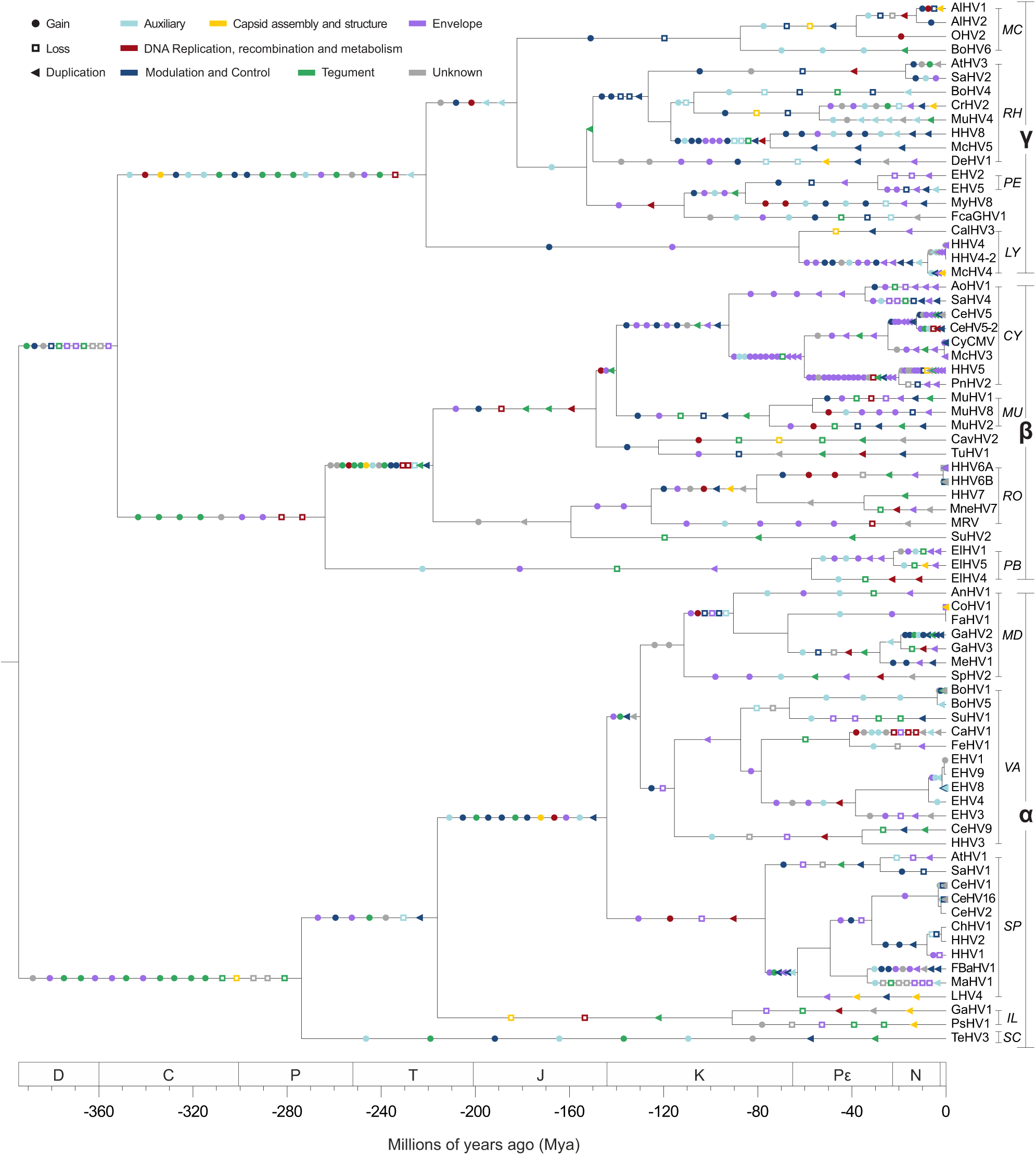
The evolution of the domain repertoire of the family *Herpesviridae.* Using the same tree shown in Figure 1, events of domain gain, loss and duplication (in this order) were mapped along the branches where they most likely took place according to their ancestral character reconstruction. The colour scheme shown here is the same used in Figure 2.

To assess the importance of each of these events over time, and to explain how domains of distinct functions evolved, Figure 5 shows the distribution of domain gains, losses and duplications across four time intervals, starting from the tMRCA (time to MRCA) of all herpesviruses.

**Figure 5.**
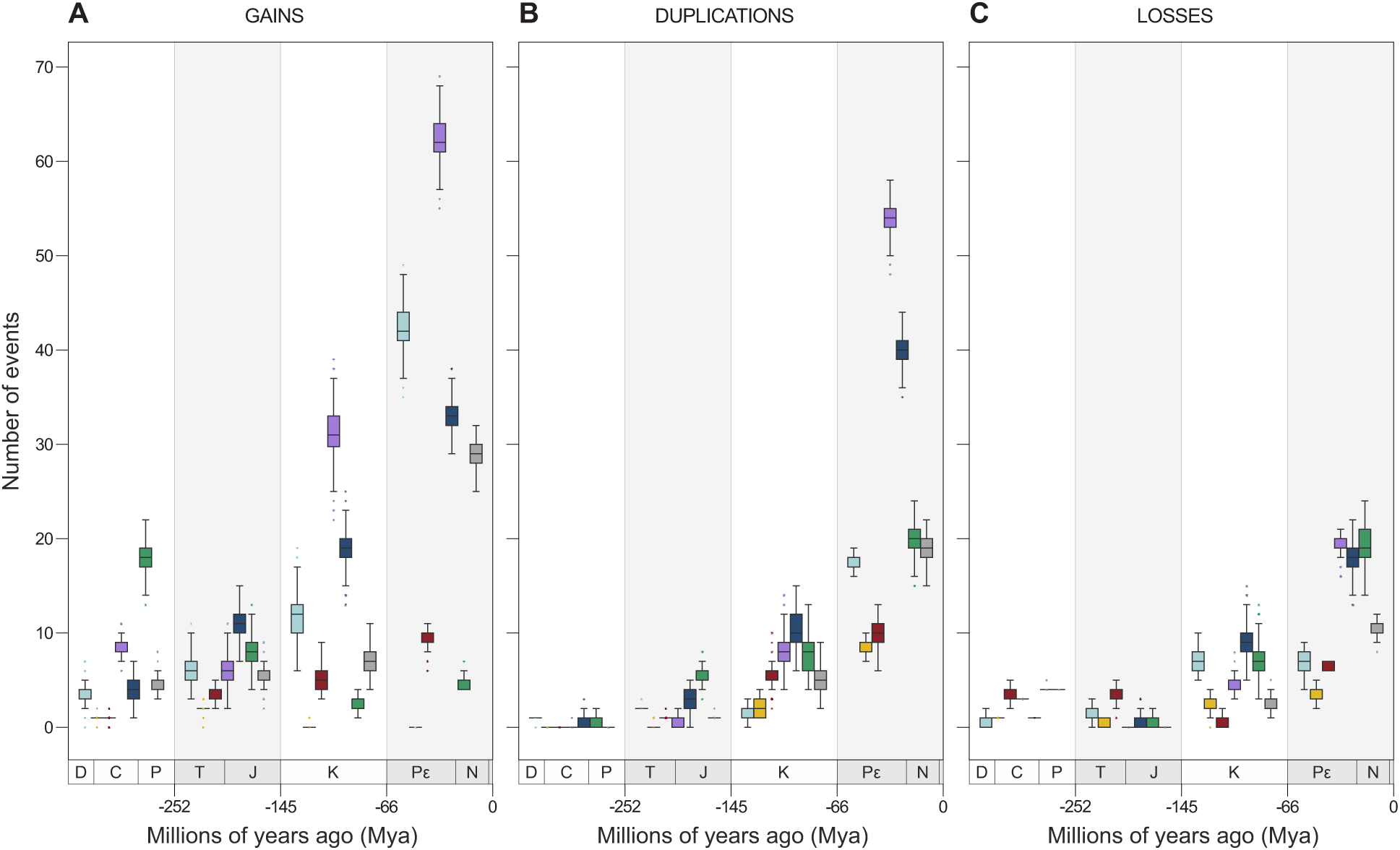
Frequency of gains, losses and duplications of domains and molecular functions across four time intervals. By performing 1000 simulations, each event had its time of occurrence sampled randomly along the branch it belongs to, allowing the events shown in Figure 4 to be split highlighting their frequency in four chronological bins: Devonian-Carboniferous-Permian (D-C-P); Triassic-Jurassic (T-J); Cretaceous (K); and Paleogene-Neogene-Quaternary (P_ε_-N-Q). In this way, the scenario observed here summarizes the one shown in Figure 1. Since the number of branches increases exponentially, absolute values at each bin in the subplots of Figure 5 cannot be directly compared due to differences in scale. Therefore, the purpose of these plots is to highlight what events and domain functions were predominant at each interval.

It is possible to observe, for example, that events of domain gain outnumber losses and duplications in all intervals. Between the Devonian (D) and the end of the Cretaceous (K), similar numbers of losses and duplications of domains of distinct functions were observed. As the number of sampled taxa differs in Alpha-, Beta- and GammaHVs, to compare the impact of domain gains, losses and duplications in the evolution of their repertoires, we established event rates by normalizing the total number of events (e) by the tMRCA of *Hespesviridae* (∼394 Mya), and the total number of taxa (spp) per subfamily (e x tMRCA^−1^/spp) (Figure 6A).

**Figure 6.**
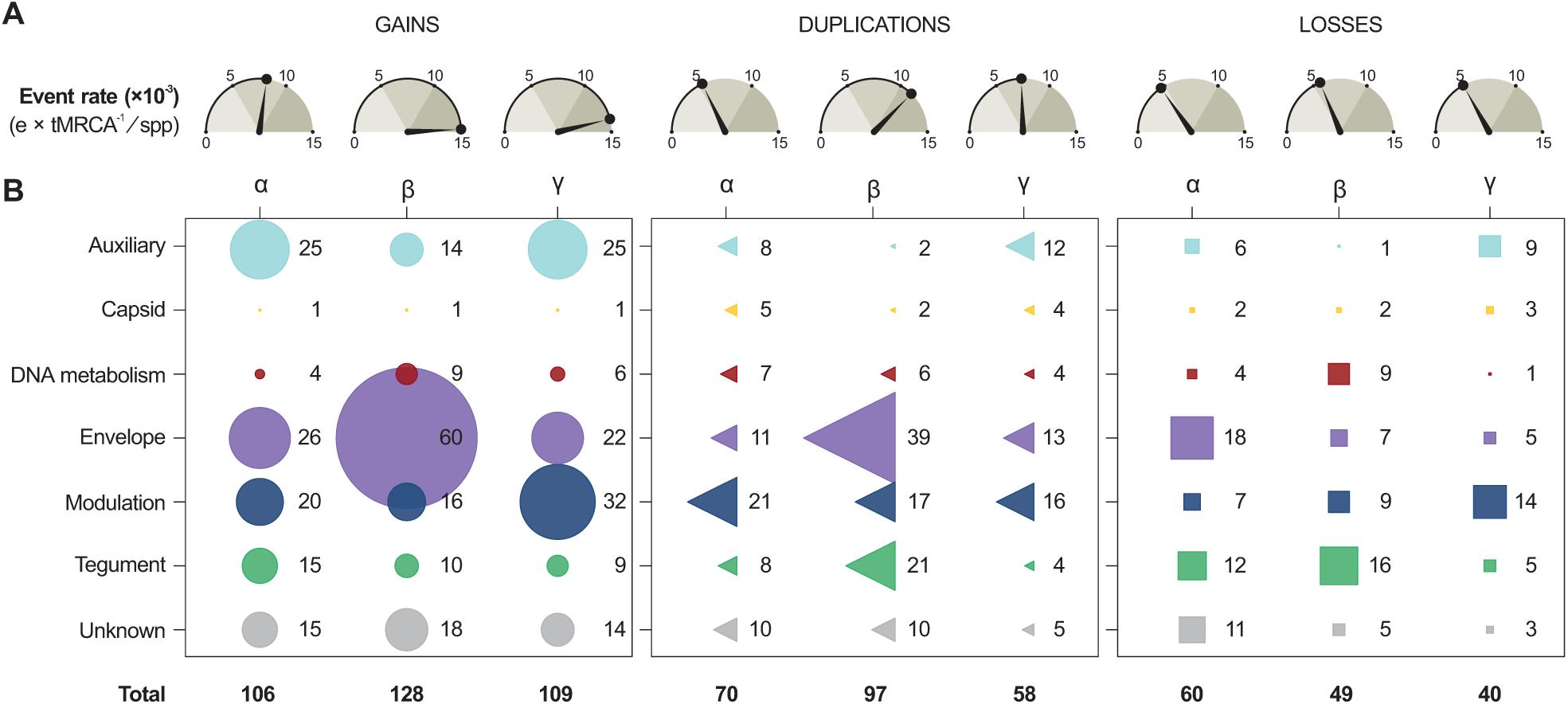
The impact of gains, losses and duplications shaping the domain repertoires of HV subfamilies. A) Number of events (× 10^−3^) normalized by tMRCA (in Myr) and number of taxa (spp). B) Absolute frequency of gains, losses and duplication of domains from distinct functional categories.

For domain gains, these results revealed that Betaand GammaHVs evolved under higher rates of acquisitions (14.77 and 13.83 x 10^−3^ gains per Myr per species), nearly twice as high as AlphaHVs (8.15 x 10^−3^). The levels of domain losses in all subfamilies were very similar (around 5 x 10^−3^), but lower than gains and duplications. Finally, rates of domain duplications varied among the three HV groups, with BetaHVs showing the highest rate (11.19 x 10^−3^ duplications per Myr per species). Overall, rates of events expanding the repertoire of *Hespesviridae* (gains + duplications) were nearly four times higher (19.22 x 10^−3^) than those causing repertoire contraction (domain losses, 5.04 x 10^−3^).

### The primordial domain repertoire of HVs

Prior to the split of HVs into their three known subfamilies, the last common ancestor of all herpesviruses (∼394 Mya) encoded a minimal set of 41 domains (Table 1). In this set, nearly one third of the domains (27 out of 41) are involved in processes of ‘Capsid assembly and structure’ or ‘DNA Replication, recombination and metabolism’, an aspect that matches previous analysis of functional conservation in HVs (Alba et al. 2001). Domains encoded by 33 HV core genes are found in this conserved set, which are shown in Table 1 with their corresponding annotation in the HHV1 genome. The following sections provide details about how these events impacted the evolution of HV domain repertoires and each functional category.

**Table 1.**
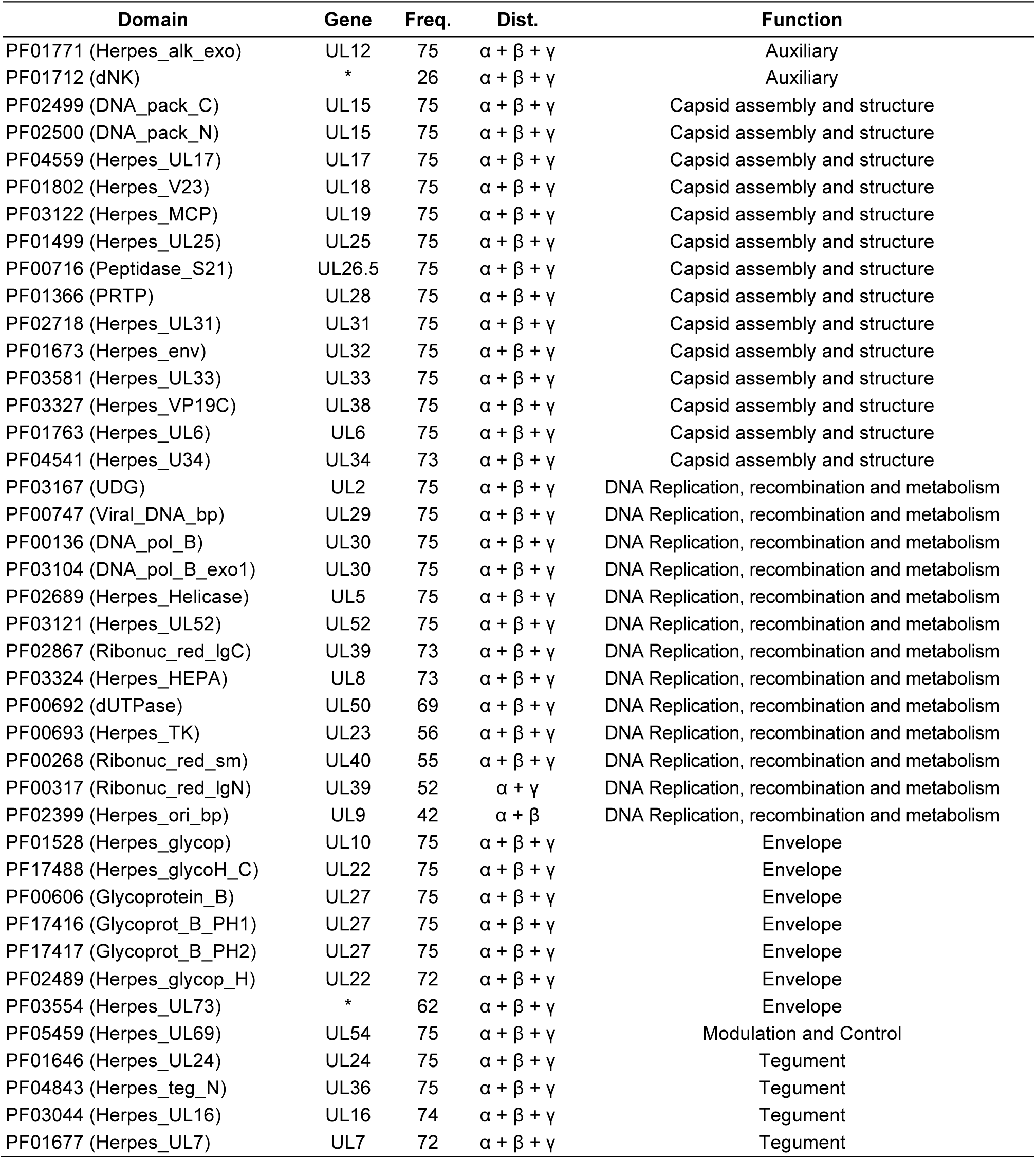
Minimal set of 41 domains encoded by the most recent common ancestor of herpesviruses. The domains are shown with their absolute frequency (Freq.) among the fully sequenced HV (75 = shared by all HVs), their distribution (Dist.) among the three subfamilies (a, p and Y), and their functional categories. Genes are named as found in the HHV1 genome annotation. * = Genes with no homologs in HHV1.

| Domain | Gene | Freq. | Dist. | Function |
| --- | --- | --- | --- | --- |
| PF01771 (Herpes_alk_exo) | UL12 | 75 | $\alpha + \beta + \gamma$ | Auxiliary |
| PF01712 (dNK) | * | 26 | $\alpha + \beta + \gamma$ | Auxiliary |
| PF02499 (DNA_pack_C) | UL15 | 75 | $\alpha + \beta + \gamma$ | Capsid assembly and structure |
| PF02500 (DNA_pack_N) | UL15 | 75 | $\alpha + \beta + \gamma$ | Capsid assembly and structure |
| PF04559 (Herpes_UL17) | UL17 | 75 | $\alpha + \beta + \gamma$ | Capsid assembly and structure |
| PF01802 (Herpes_V23) | UL18 | 75 | $\alpha + \beta + \gamma$ | Capsid assembly and structure |
| PF03122 (Herpes_MCP) | UL19 | 75 | $\alpha + \beta + \gamma$ | Capsid assembly and structure |
| PF01499 (Herpes_UL25) | UL25 | 75 | $\alpha + \beta + \gamma$ | Capsid assembly and structure |
| PF00716 (Peptidase_S21) | UL26.5 | 75 | $\alpha + \beta + \gamma$ | Capsid assembly and structure |
| PF01366 (PRTP) | UL28 | 75 | $\alpha + \beta + \gamma$ | Capsid assembly and structure |
| PF02718 (Herpes_UL31) | UL31 | 75 | $\alpha + \beta + \gamma$ | Capsid assembly and structure |
| PF01673 (Herpes_env) | UL32 | 75 | $\alpha + \beta + \gamma$ | Capsid assembly and structure |
| PF03581 (Herpes_UL33) | UL33 | 75 | $\alpha + \beta + \gamma$ | Capsid assembly and structure |
| PF03327 (Herpes_VP19C) | UL38 | 75 | $\alpha + \beta + \gamma$ | Capsid assembly and structure |
| PF01763 (Herpes_UL6) | UL6 | 75 | $\alpha + \beta + \gamma$ | Capsid assembly and structure |
| PF04541 (Herpes_U34) | UL34 | 73 | $\alpha + \beta + \gamma$ | Capsid assembly and structure |
| PF03167 (UDG) | UL2 | 75 | $\alpha + \beta + \gamma$ | DNA Replication, recombination and metabolism |
| PF00747 (Viral_DNA_bp) | UL29 | 75 | $\alpha + \beta + \gamma$ | DNA Replication, recombination and metabolism |
| PF00136 (DNA_pol_B) | UL30 | 75 | $\alpha + \beta + \gamma$ | DNA Replication, recombination and metabolism |
| PF03104 (DNA_pol_B_exo1) | UL30 | 75 | $\alpha + \beta + \gamma$ | DNA Replication, recombination and metabolism |
| PF02689 (Herpes_Helicase) | UL5 | 75 | $\alpha + \beta + \gamma$ | DNA Replication, recombination and metabolism |
| PF03121 (Herpes_UL52) | UL52 | 75 | $\alpha + \beta + \gamma$ | DNA Replication, recombination and metabolism |
| PF02867 (Ribonuc_red_lgC) | UL39 | 73 | $\alpha + \beta + \gamma$ | DNA Replication, recombination and metabolism |
| PF03324 (Herpes_HEPA) | UL8 | 73 | $\alpha + \beta + \gamma$ | DNA Replication, recombination and metabolism |
| PF00692 (dUTPase) | UL50 | 69 | $\alpha + \beta + \gamma$ | DNA Replication, recombination and metabolism |
| PF00693 (Herpes_TK) | UL23 | 56 | $\alpha + \beta + \gamma$ | DNA Replication, recombination and metabolism |
| PF00268 (Ribonuc_red_sm) | UL40 | 55 | $\alpha + \beta + \gamma$ | DNA Replication, recombination and metabolism |
| PF00317 (Ribonuc_red_lgN) | UL39 | 52 | $\alpha + \gamma$ | DNA Replication, recombination and metabolism |
| PF02399 (Herpes_ori_bp) | UL9 | 42 | $\alpha + \beta$ | DNA Replication, recombination and metabolism |
| PF01528 (Herpes_glycop) | UL10 | 75 | $\alpha + \beta + \gamma$ | Envelope |
| PF17488 (Herpes_glycoH_C) | UL22 | 75 | $\alpha + \beta + \gamma$ | Envelope |
| PF00606 (Glycoprotein_B) | UL27 | 75 | $\alpha + \beta + \gamma$ | Envelope |
| PF17416 (Glycoprot_B_PH1) | UL27 | 75 | $\alpha + \beta + \gamma$ | Envelope |
| PF17417 (Glycoprot_B_PH2) | UL27 | 75 | $\alpha + \beta + \gamma$ | Envelope |
| PF02489 (Herpes_glycop_H) | UL22 | 72 | $\alpha + \beta + \gamma$ | Envelope |
| PF03554 (Herpes_UL73) | * | 62 | $\alpha + \beta + \gamma$ | Envelope |
| PF05459 (Herpes_UL69) | UL54 | 75 | $\alpha + \beta + \gamma$ | Modulation and Control |
| PF01646 (Herpes_UL24) | UL24 | 75 | $\alpha + \beta + \gamma$ | Tegument |
| PF04843 (Herpes_teg_N) | UL36 | 75 | $\alpha + \beta + \gamma$ | Tegument |
| PF03044 (Herpes_UL16) | UL16 | 74 | $\alpha + \beta + \gamma$ | Tegument |
| PF01677 (Herpes_UL7) | UL7 | 72 | $\alpha + \beta + \gamma$ | Tegument |

### Domains with auxiliary functions

Domains classified as ‘Auxiliary’ perform distinct enzymatic and accessory functions, and were mainly acquired by Alpha- and GammaHVs, between the Paleogene (Pe) and the Quaternary (Q) periods (Figure 5A). In these periods, AlphaHVs acquired five auxiliary domains more than one time, they are: Lipase; dNK; Herpes_IR6; zf-RING_UBOX; and Keratin_B2_2. For GammaHVs, three domains were gained in multiple independent events: Lectin_C; DHFR_1; and RINGv.

Duplications of such domains were not as common as in other functional categories, being kept at very low levels until the Cretaceous (Figure 5A). Only from the Paleogene a few domains got duplicated, especially in GammaHVs, which expanded the number of copies of the domains Herpes_LP and Herpes_ORF11. At the same period, AlphaHVs experienced multiple duplications of Herpes_IR6, a domain only found in this subfamily.

Losses were uncommon among ‘Auxiliary’ domains, however, a few of them were lost in more than one occasion, in distinct time intervals. The domain dNK (Deoxynucleoside kinase), for example, was lost in most Alpha- and BetaHVs along the Triassic-Jurrassic interval. Later, between the Paleogene and Quaternary periods, AlphaHVs from the genus *Simplexvirus* lost zf-RING_2 (a Ring finger domain) at least three times. Finally, at the same period, GammaHVs from different genera lost one of their multiple copies of Herpes_ORF11, a domain found in dUTPase proteins.

### Capsid domains: conservation as a rule

The functional category of domains implied in ‘Capsid assembly and structure’ were the most conserved and less flexible group. Among the 17 existing capsid domains, 14 were inherited from the common ancestors of *Herpesviridae* (Table 1), meaning that only three domains of this kind were acquired along the evolution of these viruses (Figure 5A), one per subfamily (Figure 2 and Figure 6B). AlphaHVs acquired the domain Herpes_UL35 along the Triassic-Jurassic interval; and at the same period, BetaHVs acquired the domain HV_small_capsid; and GammaHVs acquired the domain Herpes_capsid much earlier, most likely along the Cretaceous and Permian period.

Although very rare, duplications of capsid domains were slightly more common than gains. The domain Herpes_env is an interesting exception, being duplicated in most Beta- and GammaHVs, while present as single copy in AlphaHVs (Figure 2). This very domain family was also the one facing more losses, which took place especially in more recent times (PE-Q interval).

### Domains performing DNA-related functions

As observed for ‘Capsid’ domains, the functional category ‘DNA Replication, recombination and metabolism’ represents the second most conserved and less flexible set of domains of HVs. Apart from 13 domains already present in the primordial repertoire of *Herpesviridae* ancestors (Table 1), few DNA-related domains were acquired. Important exceptions to this rule are: DNAPolymera_Pol, a catalytic subunit independently acquired by *Mardivirus* and *Simplexvirus*, and; Herpes_DNAp_acc, a DNA replication accessory factor gained by ancestors of GammaHVs. Some domains were most likely acquired from other viruses by Horizontal Gene Transfer (HGT), such as: (i) Parvo_NS1, a domain encoded by Parvovirus non-structural protein NS1 homologs, involved in viral genome replication (G0:0019079), found specifically in HHV6 and MuHV2; (ii) YqaJ, a viral recombinase domain independently acquired by MyHV8 and MuHV8, and; *(iii)* Rep_N, found in HHV6, but originally encoded by Adeno-associated virus (AAV), found in proteins involved in viral replication and integration.

Duplications in this functional category were mostly restricted to three domain families: Herpes_UL42, encoded by DNA polymerase processivity factors of AlphaHVs; Herpes_PAP, found in polymerase accessory proteins of BetaHVs; and more importantly, dUTPase, observed as two copies in several GammaHVs.

Losses of DNA-related domains, albeit uncommon, were recurrent mainly in BetaHVs (Figure 6B), which lost some domains inherited from the common ancestors of HVs, such as Ribonuc_red_lgN and Ribonuc_red_sm, which are ribonucleotide reductase domains; and Herpes_TK, a herpesviral thymidine kinase. These domains are mostly absent in all BetaHVs since the Permian and Triassic periods.

### Envelope domains: a highly flexible and redundant functional group

The functional category of ‘Envelope’ domains was undoubtedly the most flexible and redundant domain group in herpesviruses (Figure 2), which were especially acquired by BetaHVs of the genus *Cytomegalovirus* (Figure 6B). Overall, the gain rates of envelope domains were much higher than those of other functional categories (Figure 5A). The following domains, which are of animal origin (Holzerlandt et al. 2002), were gained in multiple and independent events, especially between the Paleogene and Quaternary periods: ig (Immunoglobulin domain); V-set (immunoglobulin V-set domain); C1-set (immunoglobulin C1-set domain); C2-set_2 (CD80-like C2-set immunoglobulin domain); MHC_I (Class I Histocompatibility antigen); TNFR_c6 (TNFR/NGFR cysteine-rich domain); 7tm_1 (7 transmembrane receptor); and 7TM_GPCR_Srsx (Serpentine type 7TM GPCR chemoreceptor Srsx).

Duplication of envelope domains were mostly common from the Paleogene to present times, being very infrequent until the end of the Jurassic period (Figure 1B). Proportionally to the rates of gains, BetaHVs experienced frequent events of envelope domain duplications (Figure 6B), especially involving those most likely acquired by HGT. For AlphaHVs, the envelope domain Herpes_glycop_D, found in glycoproteins D/GG/GX, was duplicated in at least 3 independent occasions. Duplications of envelope domains in GammaHVs took place mostly along the PE-N-Q interval, involving domains unique to this subfamily, in particular Herpes_BLLF1, encoded by a major outer envelope glycoprotein, and the domains encoded by latent membrane proteins, Herpes_LMP1 and Herpes_LMP2.

Losses of envelope domains were kept at low rates until the Late Cretaceous (Figure 5C). Notably, these losses took place predominantly among AlphaHVs (Figure 6B), which lost herpesvirus-specific domains in multiple occasions, they are: Herpes_glycop_D; Herpes_glycop_H; Herpes_UL43; Herpes_UL49_5; Herpes_UL73 and Herpes_US9.

### Domains involved in Modulation and Control

Domains encoded by proteins implied in ‘Modulation and Control’ of host physiology and gene expression evolved under a regime similar to that of ‘Envelope’ domains, with distinct subfamilies, genera, and species developing their specific repertoires. Only a single modulation domain is shared among all HVs species: Herpes_UL69, a transcriptional regulator inherited from the ancestors of all HVs (Table 1). The rates of gains, duplications, and losses of these domains increased steadily over time, being among the highest rates when compared to other functional categories (Figure 5). Acquisitions of domains in this category were mostly common among members of the subfamily *Gammaherpesvirinae* (Figure 6B), which gained at least 20 modulation domains, now strictly found in a few species, such as: the interleukin domains IL6, IL17, IL22, and IL31; the interferon regulatory factors IRF and IRF-3; the cyclin domains K-cyclin_vir_C and Herp-Cyclin; and other domains participating in distinct pathways, such as CARD; BH4; Bcl-2_3; bZIP_2; TNF; EBV-NA3; Herpes_LAMP2; and KSHV_K8. Finally, two interleukin domains - IL8 and IL10 -, gained at multiple independent events, are now present in all subfamilies, although not in all species (Figure 2).

Duplications of modulation domains were frequent all over the evolution of HVs, and most of them involved domains specific to a single HV subfamily. AlphaHVs, for example, obtained duplicates that are now ubiquitous among them, as the Zinc finger domains zf-C3HC4, zf-C3HC4_2, and zf-C3HC4_3; and the viral regulatory domains Herpes_IE68, and Herpes_ICP4_C. BetaHVs, on the other hand, obtained extra copies of domains found only in a few viral lineages, as the immediate early protein domains HHV6-IE (unique to *Roseolovirus)* and Herpes_IE1 (unique to *Cytomegalovirus).* In GammaHVs, the main domains with duplicates were: Sushi domain, mostly found in multiple copies in *Rhadinovirus,* and; the death effector domain (DED), mainly encoded by members of *Rhadinovirus* and *Percavirus.*

Modulation domains were among the main functions lost after the Jurassic period (Figure 5C), and a large number of them involved GammaHVs (Figure 1B). The oldest losses of modulation domains in this subfamily took place most likely in the Cretaceous period (K), when the domain Bcl-2, an apoptosis regulator, was independently extinct at ancestors of *Macavirus, Rhadinovirus,* and *Percavirus.* In BetaHVs, the interleukin domain IL8 was lost at least four times, mostly along the P_ε_-N-Q interval, similar to what happened with AlphaHVs, where the zinc finger domain zf-C3HC4_2 was extinct in distinct species of *Simplexvirus* and *Mardivirus.*

### Domains of tegument proteins

Tegument proteins establish mutual interactions forming a tight matrix that links the capsid and the envelope of HVs (Mocarski 2007; Owen et al. 2015). ‘Tegument’ domains are highly conserved across subfamilies, and four of them were inherited from the common ancestors of HVs: Herpes_UL7; Herpes_UL16; Herpes_UL24; and Herpes_teg_N, the last two still present in all HV species (Figure 2). The total numbers of tegument domains acquired by each subfamily were similar, and notably this was the only functional category where rates of domain gain decreased over time (Figure 5A), with the level of acquisition reaching its peak in early times, between the Devonian and the Permian. Along these periods, AlphaHVs acquired a set of seven domains still encoded by almost all their members: Herpes_UL14; Herpes_UL21; Herpes_UL36; Herpes_UL37_1; Herpes_UL46; Herpes_UL51; and Pkinase. At the same time interval, BetaHVs acquired four subfamily-specific domains: Herpes_U59; Herpes_pp85; DUF2664; and UL97, a domain with protein kinase activity (GO:0004672). Finally, during those early times, GammaHVs acquired their own tegument domains, such as: Herpes_BTRF1; Herpes_BLRF2; Tegument_dsDNA; DUF832; DUF848; and DUF2733.

Following an inversely proportional trend of domain gains, the rate of ‘Tegument’ domain duplication increased over time (Figure 5B), with BetaHVs providing a major contribution to that (Figure 6B). The main duplications in this subfamily took place in the Pε-N-Q interval, and involved the domains Herpes_pp85, Herpes_UL82_83, and US22. Finally, duplications in Alpha- and GammaHVs involved mostly the domains Pkinase and Tegument_dsDNA, respectively.

Overall, the functional category ‘Tegument’ experienced the largest number of losses (Figure 5B and Figure 6B), being the main function lost between the Paleogene and Quaternary, especially by BetaHVs. In this period, this viral group lost copies of US22, while a few domains were extinct in some species, in particular the domain U71, mostly extinct in *Cytomegalovirus;* Herpes_U30, lost in some *Proboscivirus;* and DUF2664, independently lost in distinct genera.

### Domains with other/unknown functions in HVs

Although domains classified with other/unknown functions do not make a functional category per se, it is important to highlight a few instances of gains, duplications and losses of such domains, as it could provide hints on their possible role on HV infection cycle. Overall, although at low levels, the flux of such domains in and out of HV repertoires was constant. Between the Devonian and the Permian periods, AlphaHVs gained domains that now are ubiquitous in their subfamily, such as Herpes_UL4 and Herpes_UL3 (found in nuclear proteins), US2, and Gene66. Around the same periods, BetaHVs acquired a set of domains to date found only among them: Herpes_U5; Herpes_U55; DUF587; and DUF570. GammaHVs also gained their subfamily-specific domains, in particular Herpes_BBRF1 and DUF717, now present in most members of that group. Finally, most probably in the Devonian, ancestors of Beta- and GammaHVs acquired three domains still conserved in all their species: Herpes_UL87; Herpes_UL92; and Herpes_UL95.

Duplications were mainly observed in recent times (P_ϵ_-N-Q) (Figure 5B), with AlphaHVs obtaining duplicates of Gene66 and Collagen, the later also duplicated in a few GammaHV species. In BetaHVs of the genus *Roseolovirus,* multiple duplications of DUF865 took place, and now such viruses encode from 8 to 22 copies of this uncharacterized domain. Finally, losses involved mostly domains unique to AlphaHVs (Figure 6B), such as Herpes_UL3, Gene66, and US2, which were lost along the Neogene and Paleogene periods.

## DISCUSSION

To our knowledge, this study is the first to provide a thorough overview on the evolution of domain repertoires of herpesviruses, tracking gains, losses, and duplications of domains from distinct functions along the viral phylogeny. Since all possible ORFs were scanned for domains disregarding their sizes, initial codons, or whether or not they were annotated in the original genome annotations, our approach detected a large set of at least 274 domains in HVs. It was only possible due to the sensitivity of HMM profile screenings, which allows the detection of remote homology between sequences, a task that conventional methods of sequence comparison are unable to achieve (Mistry et al. 2013). The domain repertoires of herpesviruses evolved at distinct rates, and the interplay among gains, losses and duplications of domains from distinct functional categories determined the current composition of HV domain repertoires.

The evolution of HVs was largely characterized by high rates of domain acquisition. Since the early Devonian herpesviruses nearly doubled the size of their domain repertoires, increasing from around 41 domains encoded by their ancestors, to an average of 75.6 ± 8.4 domains encoded by present-day HVs. Most domains acquired by these viruses are from the functional categories ‘Auxiliary’, ‘Envelope’ and ‘Modulation and Control’, and were mainly gained after the Jurassic (Figure 5A), coinciding with the rapid adaptive radiation underwent by mammalian species in that period (Lee and Beck 2015). As host range and cell tropism are mostly defined by cell surface receptors, intracellular factors, and the ability of viruses to manipulate host physiology (Werden et al. 2008; Grinde 2013), the high levels of gains of domains performing ‘Auxiliary’, ‘Envelope’ and ‘Modulation and Control’ functions was probably an adaptive strategy adopted at least by mammalian HVs to cope with the rapid diversification of their host species. Many of these domains allowed viruses to evade and modulate the host immune system (Bernet et al. 2003), particularly by mimicking and/or blocking cellular processes, giving HVs the ability to establish latency and lifelong infections (Holzerlandt et al. 2002; Grinde 2013).

Duplications were particularly common for domains involved in ‘Tegument’, ‘Modulation and Control’ and ‘Envelope’ functions. The rates of these events varied across subfamilies, with BetaHVs showing the highest rate of duplications. In this HV subfamily, ‘Envelope’ domains were the main category with duplicates, most of them involving cytomegaloviruses. When a gene or domain gets duplicated, their copies can have two possible fates: domain loss, or neofunctionalization (Wagner 1998). Especially in Beta- and GammaHVs, the overall rates of domain duplications were higher than losses (Figure 6), indicating that most duplicated domains were kept in their repertoires. With duplicates following independent evolutionary paths, neofunctionalization can operate by means of multiple mutations leading to radical changes in function, or by fine changes on protein affinity and specificity (Levasseur and Pontarotti 2011). Several Beta- and GammaHV domain duplicates are known to have diverged to perform distinct functions. This is observed, for example, in dUTPase homologs, where neofunctionalization generated a broad set of proteins that perform distinct roles in viral infection (Davison and Stow 2005; McGeoch et al. 2006). Another known example of duplication leading to neofunctionalization involves the domain Herpes_glycop_D, found only in AlphaHVs and present as two copies in most species: the glycoprotein D (gD), an envelope protein that inhibits cell-to-cell adhesion and plays a role in cell entry and fusion (Zhang et al. 2011); and the glycoprotein G (gG), which by mimicry shows chemokine binding activity, antagonizing cellular chemokine receptors (Bryant et al. 2003; Van de Walle et al. 2009).

Nearly a quarter of the domains (67 out of 274) in the repertoire of *Herpesviridae* are currently present with more the one copy in some species. As retention of domain duplicates were higher than losses in most HVs, our results indicate that, apart from the aforementioned examples, neofunctionalization may be a more common process in HVs than previously thought. Cases of duplication illustrated in Figure 2 provide room for further investigations about the functions carried out by duplicates during infections.

The rates of domain losses for each HV subfamily were very similar, although the functional categories of lost domains differed across these viral groups (Figure 6). Although domain gains and duplications outnumbered losses, the elimination of domains from HV repertoires played a central role as a source of genetic variation across species, genera, and subfamilies. Along the evolutionary history, gene losses can be either an adaptive or neutral phenomenon, which brings about two main hypotheses explaining the evolution of genomes: the ‘less-is-more’ hypothesis, in a context of adaptive evolution, and; the ‘reductive evolution’ hypothesis, in a scenario of neutrality (Albalat and Canestro 2016). Considering the less-is-more hypothesis, while HVs evolve and explore new hosts or tissues, adaptations in response to environmental changes and new pressures are essential to maintain viral fitness (Daugherty and Malik 2012; Grinde 2013). If a domain proves to be highly antigenic in a certain biological context, instead of evading the host defence via multiple mutations, losses of whole domains can be an efficient and fast mode of adaptation (Daugherty and Malik 2012; Elde et al. 2012; Albalat and Canestro 2016). Another important aspect of adaptive evolution via domain losses regards the negative stoichiometric effect caused by the excessive expression of certain gene products, a phenomenon known as ‘dosage sensitivity’ (Albalat and Canestro 2016). Interestingly, it was found that some eukaryote gene families associated with ‘DNA-related’ functions tend to be duplication-resistant, and their extra copies are prone to be lost (Albalat and Canestro 2016). Our results have revealed similar trends for HV domains associated with ‘DNA Replication, recombination and metabolism’, which have shown low rates of gains and duplications (see Figure 2, Figure 5C, and Figure 6B), suggesting that domains belonging to this functional category, when duplicated, have a low probability of retention, being quickly purged from viral populations as a result of the deleterious effects of dosage imbalance (Levasseur and Pontarotti 2011; Albalat and Canestro 2016).

Apart from dosage sensitivity, another aspect favouring the loss of duplicates is the ‘dominant negative effect’, a phenomenon that emerges when a second copy of a protein interferes with the functioning of its original version (Levasseur and Pontarotti 2011). Such mechanism can lead, for example, to the formation of defective complexes (Perica et al. 2012), and as such, it could be a possible explanation for the low frequency of duplicates in domains involved in ‘Capsid assembly and structure’ (see Figure 2, Figure 5C, and Figure 6B). Probably, keeping duplicates of such protein domains may lead to structurally distinct copies of capsid subunits, which could potentially harm the capsid structural integrity by dominant negative subunits.

In the evolution of ‘Tegument’ domains, losses were very common, while their gains exceptionally decreased over time. Tegument proteins are known to establish a myriad of interactions among each other in the space between the capsid and the viral envelope, strategic position that allow them to perform structural and/or modulatory roles upon cell entry (Owen et al. 2015). Inside mature virions, these proteins create a densely packed and symmetric inner layer of proteins surrounding the capsid, and an outer layer of loosely packed proteins around it, closer to the viral envelope (Zhou et al. 1999; Yu et al. 2011). Given such heterogeneous structural and functional properties, it is not clear what phenomenon could have played a major role driving losses and preventing gains of tegument domains. Given their essential roles in viral infection, and their poor levels of conservation across HV subfamilies (Figure 2), further studies would be necessary to explain why these viral groups now encode their own specific sets of tegument domains.

Finally, considering the hypothesis of regressive evolution in a context of neutrality, genetic drift can also operate removing domains according to their levels of conditional dispensability. A genetic element can be considered dispensable when the environmental pressures justifying their maintenance in the genome cease to exist, generating a condition where keeping the gene/domain is not strictly advantageous (Albalat and Canestro 2016). This phenomenon was experimentally demonstrated, for example, in Poxviruses, where losses of duplicates were frequent upon absence of specific selective pressures (Elde et al. 2012). In scenarios of domain neofunctionalization after duplication, one of the copies can keep its original function, while the other can evolve faster. Most extra copies of genes/domains tend to accumulate loss-of-function mutations leading to the generation of pseudogenes, which can be lost due to their high levels of dispensability (Wagner 1998). In a similar way, domains that are functionally linked in pathways/complexes with domains encoded by pseudogenes can be co-eliminated in consequence of functional bias (Albalat and Canestro 2016). Since our search for domains screened all possible ORFs, disregarding their sizes or initial codons, we could not ascertain whether or not all domains here reported belong to functional genes. Further experimental studies would be necessary to clarify this aspect, and to determine what mode of evolution (adaptive or neutral) prevails at shaping the evolution of HV repertoires and driving losses of domains from distinct functional categories.

The evolutionary origins of viruses remain unclear despite decades of debates, raising distinct (and to some extent opposing) hypothesis to explain their origins, such as: *(i)* the ‘Escape’ hypothesis, which considers viruses as genetic elements that escaped from cells and expanded themselves by gene acquisitions; *(ii)* the ‘Regressive’ hypothesis, which claims that viruses derive from free-living unicellular organisms that lost several genes, thus becoming obligate intracellular parasites; and (*iii*) the ‘virus-first’ hypothesis, which asserts that viruses originated before cellular organisms, and co-evolved with them (Wessner 2010; Nasir and Caetano-Anolles 2015; Harish et al. 2016).

Using phylogenetic and domain composition information, our reconstruction of ancestral characters revealed that ancestors of herpesviruses encoded a primordial set of at least 41 domains, which perform fundamental functions such as ‘Capsid assembly and structure’ and ‘DNA Replication, recombination and metabolism’. As only 28 of these domains are still encoded by all HVs, at least 13 domains were lost along their evolution, proving not to be strictly essential. Despite that, events of domain accretion outnumbered losses, and HV genomes expanded progressively by capturing or generating domains *de novo* to perform specific functions (Davison 2002; Holzerlandt et al. 2002). This observation agrees mostly with the ‘Escape’ hypothesis, once acquisition of extrinsic genetic elements was predominant in herpesviruses. The family *Herpesviridae* is related to other two families of viruses also classified in the order *Herpesvirales,* they are: *Alloherpesviridae,* which have fishes and frogs as hosts, and eight fully sequenced genomes; and *Malacoherpesviridae*, which have bivalves as hosts, and two fully sequenced genomes (Davison et al. 2009). When their genomes are scanned for domains using the same approach we used for HVs, we observed that only one domain is conserved in all members of the *Herpesvirales* order: the domain DNA_pol_B, found in DNA polymerase family B. Remarkably, this domain is also encoded by tailed bacteriophages (order *Caudovirales)* (Petrov and Karam 2004), which also share a common DNA packing machinery with members of *Herpesvirales* (Rixon and Schmid 2014). Based on these similarities it has been hypothesized that these two viral orders may have had a common ancestor (Rixon and Schmid 2014), which most probably existed prior to the split between Eukaryotes and Akaryotes (Bacteria and Archaea) at least 2000 Mya, in the Paleoproterozoic Era (Lovisolo et al. 2003; Gradstein et al. 2012; Rixon and Schmid 2014). Considering these genetic and functional similarities between members of *Herpesvirales* and *Caudovirales*, it is plausible to suggest that from a small set of core genes, these DNA viruses co-evolved with their hosts, developing specific repertoires by acquiring new genes via HGT (Lovisolo et al. 2003; de Andrade Zanotto and Krakauer 2008; Krupovic and Koonin 2017) and by innovative and diversifying processes of duplication and *de novo* generation of domains (Davison 2002; Mocarski 2007).

By associating information about the domain content of large DNA viruses, and the viral phylogenetic history, ancestral character reconstruction can provide important insights about the origins of these genetic elements, and of viruses per se. Acquisitions, duplications and losses of domains over time shaped HV domain repertoires, and are important drivers of genetic diversity. The flow of domain coding regions in and out of HV genomes varied greatly in distinct time periods, and across viral groups, not just in terms of event rates, but also on domain functional groups. Remarkably, rates of domain loss were nearly four times lower than gains and duplications, an aspect that explains the expansion of HV genomes. Despite sharing a core set of 28 domains, along more than 400 millions of years of evolution, co-infections with other viruses, and co-evolution with their hosts, herpesviruses developed very specific domain repertoires at the subfamily, genus and species levels. Further studies focused on the role of such group-specific domains could help us understand which genetic elements define HV host range and tissue tropism, and could also suggest new targets for diagnosis and treatment of herpesvirus infections.

## METHODS

### Herpesvirus species and protein domain identification

To identify domains encoded by herpesvirus genomes, all possible ORFs from 75 fully sequenced genomes (Supplemental Table S1) available on NCBI Viral Genomes (Brister et al. 2015) were translated, and included in our query database. Using the program *hmmscan* implemented in HMMER-3.1 (Mistry et al. 2013), all peptides were searched against profile HMMs generated using Pfam-A seed alignments (version 31.0) (Finn et al. 2014). To decide which hits were reliable enough to be kept as true domains, strict inclusion thresholds of 0.001 were defined for both per-sequence e-value and per-domain conditional e-value. Considering these thresholds, python scripts were used to retrieve the domain repertoire of each herpesviral species from the *hmmscan* outputs. By integrating information from Pfam (Finn et al. 2014), Interpro, Pfam2GO (Mitchell et al. 2015) and previous studies (Davison 2002; Mocarski 2007), each domain was classified by its main functional category, such as: ‘Auxiliary’; ‘Capsid assembly and structure’; ‘DNA Replication, recombination and metabolism’; ‘Envelope’; ‘Tegument’; ‘Modulation and Control’; or ‘Other/Unknown’ in case it does not fit in any of the previous categories.

### Phylogenetic analysis

To understand the evolution of protein domains, the inference of a herpesviral phylogenetic tree is an essential step. Using MAFFT (Katoh and Standley 2013) we generated multiple sequence alignments (MSAs) of conserved viral proteins encoded by UL15, UL27 and UL30 genes, and amino acid substitution models were determined using ProtTest (Darriba et al. 2011). The MSAs were used as partitions on *Beast to reconstruct a time-calibrated maximum clade credibility (MCC) tree using a Markov Chain Monte Carlo (MCMC) Bayesian approach implemented in BEAST v2.4.5 (Bouckaert et al. 2014). Two divergence time priors were used to date internal nodes: the *Herpesviridae* MRCA (most recent common ancestor), according to (McGeoch and Gatherer 2005); and the MRCA of HHV1 and HHV2 according to (Wertheim et al. 2014). Priors of monophyletic constrains were incorporated following taxonomic classification provided by ICTV (King et al. 2018). The analysis was run for 35 million generations using the Yule model as a coalescent prior, and relaxed (uncorrelated lognormal) molecular clock.

### Ancestral state reconstruction

Taking the domain repertoires of the existing herpesvirus species, a matrix of domain counts was constructed using Python scripts. Applying a parsimony reconstruction method implemented in Mesquite (Maddison and Maddison 2009), this matrix was used to reconstruct the ancestral states of each domain. This reconstruction allowed us to infer events of domain gain, losses and duplications, and to map them along the branches of the HV phylogeny using iTOL (Letunic and Bork 2016), highlighting a possible parsimonious scenario for the evolution of HV domain repertoires.

## DATA ACCESS

All accession numbers of genomes, domains and GO terms mentioned in this study are listed in the Supplemental Tables.

## ACKNOWLEDGMENTS

AFB is funded by *Ciência sem Fronteiras,* a scholarship programme managed by the Brazilian federal government (CAPES, Ministry of Education, Grant number: 11911-13-1). JWP is supported by a University Research Fellowship from the Royal Society.

Authors contributions: AFP and JWP conceived and designed the study; analyzed the data, and wrote the manuscript.

